# TARP ɣ-8 is a target of ethanol that regulates self-administration and relapse in mice

**DOI:** 10.1101/2025.03.27.645788

**Authors:** Sara Faccidomo, Jessica L. Hoffman, Julie Lee, Ciarra M. Whindleton, Michelle Kim, Seth M. Taylor, April Kim, Caroline Richter, Hanna L. Seiters, Jacob M. Bryant, Ashley Chang, Evan N. Smith, Abigail E. Agoglia, Susumu Tomita, Melissa A. Herman, Clyde W. Hodge

**Author notes:** **Correspondence:** Clyde W. Hodge.

## Abstract

**Background:** Behavioral pathologies that characterize alcohol use disorder (AUD) are driven by the powerful reinforcing, or rewarding, properties of the drug. We have shown that glutamate AMPA receptor (AMPAR) activity is both necessary and sufficient for alcohol (ethanol) reinforcement. Transmembrane AMPAR regulatory protein (TARP) γ-8 is an essential auxiliary protein that regulates AMPAR expression and activity; however, the role of TARP ɣ-8 in AUD or other forms of addiction remains largely unexplored.

**Objectives:** This study investigated the mechanistic role of TARP γ-8 in operant ethanol self-administration (model of primary reinforcement) and cue-induced reinstatement of ethanol-seeking behavior (model of conditioned reinforcement) using TARP ɣ-8 heterozygous null (+/−) mice. To determine if TARP ɣ-8 signaling is targeted by ethanol use, we evaluated protein expression of TARP γ-8, GluA1, CaMKII, and PSD-95 following ethanol self-administration.

**Results:** A battery of tests evaluating food and water intake, taste reactivity, anxiety-like behavior, and object recognition memory showed no fundamental behavioral deficits in TARP γ-8 (+/−) mice, and no differences in response to acute ethanol or home-cage drinking as compared to wild-types. However, TARP γ-8 (+/−) mice exhibited significantly reduced acquisition and escalation of operant ethanol self-administration and reduced cue-induced reinstatement of ethanol-seeking behavior, with no differences in parallel sucrose-only controls. In wild-type mice, ethanol self-administration increased TARP γ-8 expression in the amygdala, nucleus accumbens, and hippocampus, and increased GluA1 expression in the amygdala and prefrontal cortex, compared to sucrose controls.

**Conclusion:** These findings highlight the specificity of TARP ɣ-8 regulation of ethanol reinforcement mechanisms and identify this crucial AMPAR auxiliary protein as a target of ethanol in reward-related brain regions, highlighting its potential for development of novel pharmacotherapies for AUD.

## Introduction

Alcohol use disorder (AUD) is a chronic, relapsing condition characterized by the progressive escalation of alcohol consumption from repeated binge/intoxication episodes to compulsive seeking and dependence. As dependence develops, the reinforcing effects of alcohol become dysregulated, driving excessive intake through maladaptive recruitment of neural circuits, including the amygdala (Koob 2021). Beyond primary reinforcement, conditioned cues associated with alcohol use are critical drivers of relapse. Clinical studies demonstrate that alcohol-paired cues promote craving and relapse during abstinence, activating reward-related regions such as the amygdala (Schneider et al. 2001). A major challenge in the field is to identify the molecular mechanisms through which alcohol alters neural systems of primary and conditioned reinforcement to sustain maladaptive alcohol seeking and relapse (Weiss and Koob 2001).

Growing preclinical evidence implicates glutamatergic neurotransmission in the reinforcing properties of alcohol (ethanol, ETOH) (Holmes et al. 2013), with particular focus on the role of AMPA receptors (AMPARs) (Hopf and Mangieri 2018). AMPARs are glutamate-gated ion channels composed of GluA1 – 4 subunits that mediate fast excitatory synaptic transmission and are essential for synaptic plasticity. Our previous work identified CaMKII-dependent AMPAR signaling in the amygdala as a key mechanism underlying the positive reinforcing effects of EtOH (Salling et al. 2016). We further demonstrated that AMPAR activation promotes escalated EtOH self-administration and cue-induced reinstatement (Cannady et al. 2013), and that EtOH self-administration increases synaptic GluA1-containing AMPAR expression in the basolateral amygdala (BLA), with GluA1 membrane trafficking required for EtOH reinforcement (Faccidomo et al. 2021). However, the upstream regulatory mechanisms controlling AMPAR function in EtOH reinforcement remain to be fully elucidated.

One critical regulatory mechanism involves transmembrane AMPAR regulatory proteins (TARPs), a family of eight auxiliary proteins (γ-1 to γ-8) that modulate AMPAR trafficking, localization, and function (Jackson and Nicoll 2011; Nicoll et al. 2006; Park et al. 2016). Among these, TARP γ-8 is of particular interest due to its restricted expression in limbic regions implicated in EtOH-related behaviors, including the prefrontal cortex (PFC), hippocampus (HPC), and BLA (Maher et al. 2016; Tomita et al. 2003). TARP γ-8 regulates AMPAR surface expression and stabilization at glutamate synapses via CaMKII phosphorylation, which promotes γ-8 binding to the C-terminus of GluA1, anchoring it within the postsynaptic density to PSD-95 (Bissen et al. 2019; Patriarchi et al. 2018). These anatomical and molecular properties position TARP γ-8 as a critical modulator of glutamate neurotransmission within EtOH-sensitive circuits that contribute to reinforcement and relapse (Koob and Volkow 2016).

Recently, we identified a novel role for TARP γ-8-bound AMPARs in mediating the reinforcing effects of EtOH, with anatomical specificity to the BLA (Hoffman et al. 2021; Hoffman et al. 2023). These findings represent the first demonstration of TARP γ-8 involvement in any addictive behavior and suggest that TARP γ-8 may serve as a pivotal regulator of EtOH-related behavioral pathology. This discovery opens a promising avenue for understanding the molecular underpinnings of AUD and developing targeted interventions.

In the present study, we investigated whether TARP γ-8 is necessary for EtOH reinforcement and relapse-like behavior using TARP γ-8 heterozygous null (γ-8 +/−) mice, which display a 50% reduction in TARP γ-8 protein expression without alterations in basal synaptic transmission (e.g., AMPA/NMDA ratio). Notably, TARP γ-8 (+/−) mice exhibit a ∼38% reduction in AMPA-evoked extrasynaptic currents in the hippocampus, suggesting that the pool of available TARP γ-8-bound AMPARs scales with TARP γ-8 expression (Rouach et al. 2005). This model provides a unique opportunity to dissect the contribution of TARP γ-8-bound AMPARs to the abuse liability of EtOH.

We hypothesized that partial deletion of TARP γ-8 would attenuate EtOH reinforcement and cue-induced relapse in a manner associated with reduced impact of EtOH on AMPAR GluA1 protein expression in key brain regions such as the amygdala. To test this, we assessed operant EtOH self-administration, relapse-like behavior and regional protein expression in TARP γ-8 (+/−) and wild-type mice, aiming to define the novel contribution of TARP γ-8 to the neural adaptations supporting EtOH use and relapse vulnerability.

## Materials and methods

### Mice

Male and female TARP γ-8 (B6; *Cacng8^tm1Ran^*/J) heterozogous null (+/−; n=26) and C57Bl/6J wildtype (+/+; n=25) mice were obtained from Jackson Labs (JAX stock #028446; Bar Harbor, ME) at 8 weeks of age. These mice were originally generated by Dr. Roger Nicoll at UCSF by targeting the TARP γ-8 locus (Cacng8) on chromosome 7 and were backcrossed to a C57BL/6 genetic background (Rouach et al. 2005). Upon arrival at UNC, all mice were genotyped by Transnetyx using real-time PCR for confirmation. One TARP γ-8 (+/−) mouse was identified upon genotyping as a (+/+) mouse and was excluded from all studies. Mice were either singly housed (cohort 1 – home cage drinking studies; n=18) or group-housed (cohort 2 & 3; operant self-administration studies; n=33) upon arrival in Techniplast cages (12” x 6.5” x 7”). Food (Isopro-3000; Granville Milling) and water were available *ad libitum* for most studies. The vivarium was maintained at 27°C and 40% humidity on a reverse 12L:12D cycle (lights off at 0700

### Appetitive Studies

Voluntary home cage food and fluid consumption studies were conducted to assess whether TARP γ-8 (+/−) mice exhibit differences in basal food and fluid consumption and preference. For the first 9 days after arrival, mice were weighed and handled daily to assess growth rate and to allow acclimation to our vivarium. Next, food and water consumption were measured in 24-hr intervals for 1 week to assess whether gross differences in fluid and food consumption were present. Fluid preference studies were conducted to assess whether these mice showed basal differences in taste reactivity to both palatable (0.03 & 0.06% Saccharin; 0.05 & 2% Sucrose) and bitter (0.015 & 0.03mM Quinine) solutions. Briefly, mice were weighed, and home cage water bottles were removed from 1000 to 1400 (starting 3 hrs into the dark cycle) and replaced with two drinking tubes containing either water or one of the sucrose/saccharin/quinine solutions. The drinking tubes consisted of a 10mL serological pipette fitted with a ball-bearing sipper tube. Mice were given access to each concentration for 2 consecutive days before moving onto the next concentration such that within a week, a preference test was conducted on two concentrations of the same solution, in ascending order of concentration. Week to week, the order of presentation of the solutions was randomized to avoid an order effect and within each week, drinking tubes alternated from left to right to avoid side preferences.

### Open Field Locomotion, Anxiety-like Behavior, and Memory

Locomotor activity, thigmotaxis, and novel object recognition were measured in Plexiglas open-field activity monitor chambers (27.9 cm^2^; ENV-510, Med Associates, Georgia, VT) as described previously (Hodge et al. 2002; Riday et al. 2012; Spanos et al. 2012; Stevenson et al. 2019). Two sets of 16 pulse-modulated infrared photobeams were located on opposite walls and recorded X–Y ambulatory movements as previously reported. Distance traveled (in cm) throughout the session was quantified by assessing the mouse’s position in the open field every 100 milliseconds. Data from each chamber were collected by a computer and were used to measure general locomotor activity and anxiolytic behavior using a light/dark insert (Med Associates) and analysis of center zone activity in the open field. Thigmotaxis was evaluated by comparing distance (cm) traveled in the center zone (inner 25% of the area) to distance traveled in the periphery (outer 75% of the area). The Novel Object Recognition Test (NORT) was conducted in a custom-made plexiglass open field arena (18” x 18” x 13”). Mice were initially habituated to the testing arena for 2, 5 min sessions. The objects used were a Lego cube (2×2×3”) and a plastic golf ball. The NORT protocol consisted of two distinct phases. After habituation, a 5 min familiarization session was conducted in which two identical objects were placed in opposite corners of the open field. Each mouse was given 5 min to freely explore the chamber and objects. Four hrs later, the NORT test was conducted. This 5 min test was identical to the familiarization session except one of the familiar objects was replaced with a novel object. The identity and location of the familiar vs novel objects were randomized across subjects. For all sessions, a camera connected to a computer running ANY-maze digital tracking software (Stoelting Co., WoodDale IL) was used to record general locomotor activity, as well as time spent with and oriented towards the zones for each object.

### Acute Ethanol Sensitivity and Blood EtOH Clearance

To assess sensitivity to the acute locomotor activating dose of EtOH, mice were habituated to an open field for 20 min, briefly removed, injected with 1.0 g/kg EtOH (20% w/v, IP) and returned to the open field for 20 min. To assess sensitivity to a sedative/hypnotic dose of EtOH, mice were injected with 4.0 g/kg EtOH (20% w/v, IP) and were placed supine in a V-shaped trough until they regained their righting reflex. Criteria to regain righting reflex was being able to right themselves 3x within 30 sec. To measure EtOH metabolism, mice were injected with 4.0 g/kg EtOH (20% w/v, IP) and tail blood was collected into a heparinized microcapillary tube at 10, 30, 60, 120, 240, and 360 min post-injection. Blood samples were analyzed using an AM1 Analox machine (Analox Instruments, Stourbridge UK).

### Home Cage Ethanol Drinking and Preference

#### 24-hr access

Mice were given 24-hr home cage access to increasing concentrations of EtOH to assess whether TARP γ-8 (+/−) mice show baseline differences in EtOH consumption and preference. Home-cage drinking was conducted as described in (Faccidomo et al. 2018; Hoffman et al. 2019; Salling et al. 2016; Stevenson et al. 2009) with the following differences. All mice had continuous access to two 50 mL ball-bearing sipper tubes containing EtOH (1-20% (w/v)) vs. water for 20 days. EtOH concentration was increased every 4 days (i.e.: days 1-4: 1%; days 5-8: 5%) Bottles were weighed, refilled, counterbalanced and rotated daily to eliminate a side preference. EtOH intake was calculated as g/kg/24-h. EtOH preference was calculated as a percentage of total fluid intake per 24-h (EtOH mls / Total mls) * 100).

#### Drinking in the Dark

Mice were given limited home cage access in the dark-cycle to assess whether TARP γ-8 (+/−) mice show differences in EtOH consumption using a modified drinking in the dark protocol. From Mon to Wed, mice were given home-cage access to 20% (v/v) EtOH via a modified 10 mL volumetric tube with ball-bearing sipper from 1000 to 1200. On Thursdays, access was increased to 4 hrs and ended at 1400 and tube readings were taken in 2 hr intervals.

### Operant Ethanol and Sucrose Self-Administration

#### Operant Conditioning Chambers

Self-administration sessions were conducted in mouse operant conditioning chambers enclosed within a sound attenuating cubicle (Med Associates, St Albans, VT) as previously described (Faccidomo et al. 2024a; Faccidomo et al. 2019; Faccidomo et al. 2015; Hodge et al. 2006; Olive et al. 2000). Each chamber contained two ultra-sensitive retractable levers (force =0.02N) located directly beneath a cue light. Responding on the “active” lever was reinforced and responding on the “inactive” lever produced no programmed consequences. Reinforcements were delivered into a drinking trough positioned adjacent to the active lever and delivery was accompanied by illumination of the cue light (800ms) and pump sound, both of which served as secondary cues. A photobeam spanned the entrance of the drinking trough to quantify head-poke entries. Lever presses made during the reinforcer delivery were measured but did not count towards the response requirement (time-out period). The chambers were connected to an interface and computer that recorded the input and behavioral output of each mouse (MED-PC for Windows v.4.1).

#### Acquisition

The acquisition protocol used in this experiment was a modified version of a previously published method (Faccidomo et al. 2009). Mice were water restricted and trained in two 16-hr overnight sessions to lever press for the delivery of a 2% sucrose (w/v) solution (0.014 ml/reinforcement). Initially, the first 25 lever presses were reinforced by a delivery of 2% sucrose (FR1). Subsequently, the response requirement increased from FR 1 to FR 2 to FR 4 in increments of 25. Starting on the third training day, operant conditioning sessions were shortened to 1-hr in duration, behavior was maintained at a FR4 schedule of reinforcement, and a modified sucrose fading procedure (Faccidomo et al. 2009; Hodge et al. 1992; Samson 1986) was implemented to facilitate EtOH self-administration and to assess genotype differences in operant responding at a variety of reinforcing solutions. All 1-hr sessions occurred in the dark phase, between 1300-1500h, 7 days/week. EtOH (v/v) was added in increasing 3% increments (3% to 30%) to the 2% sucrose solution with 3 days at each concentration. During the 34^th^ 1-hr session, the reinforcing was returned to 9% EtOH (v/v)/ 2% sucrose (w/v) for the remainder of the experiment. This is the standard reinforcing solution we use for all our self-administration studies and have published that it is a palatable solution that C57BL/6 mice consume fully (ie: consume ALL fluid in trough). The sucrose control group received 2% sucrose (w/v) only for the same number of training sessions.

#### Extinction and Cue-Induced Reinstatement

Following an extended baseline at 9% EtOH (v/v)/ 2% sucrose (w/v) or 2% sucrose (w/v) only, operant responding for EtOH or sucrose was extinguished by removing all programmed consequences that were previously associated with the lever press (reinforcer delivery, cue light, pump sound) as previously described (Cannady et al. 2013; Salling et al. 2017; Schroeder et al. 2008). Three mice were excluded from the Ethanol extinction study for the following reasons, broken inactive lever, no responding on either lever, died before this phase of the experiment. Extinction sessions occurred for 13 consecutive days. Following the 13^th^ extinction session, 3 cue-induced reinstatement sessions were conducted. During these tests, the drinking troughs were primed with 0.028mL of the reinforcing solution (volume equivalent to two reinforcers) to provide an initial gustatory stimulus at the start of the session. Tests occurred for 1 hr, during which lever pressed on the previously “active” lever, were followed by presentation of the cue light and pump sound, *without delivery of a reinforcement*.

### Gel Electrophoresis & Immunoblot

To address whether a history of EtOH or sucrose operant self-administration altered protein expression throughout the brain, mice were rapidly decapitated after 2 weeks of stable ethanol or sucrose self-administration and brains were flash-frozen in isopentane (−40°C). Thick coronal brain sections (1mm) were sliced on a cryostat (Leica Biosystems) and a circular tissue punch (1mm diameter) was bilaterally dissected from the amygdala, nucleus accumbens, dorsomedial hippocampus and medial prefrontal cortex (unilateral punch). Tissue samples were ultrasonified in a SDS homogenization buffer containing protease and phosphatase inhibitors and protein concentration was quantified using a Bradford calorimetric assay kit (BCA Assay; Pierce Biotechnology). Tissue samples (5µg) were loaded into a 4–15% Criterion TGX precast gels (Bio-Rad) with Tris Glycine-SDS running buffer (Bio-Rad) for gel electrophoresis separation. An iBlot® Semi-Dry blotting system (Thermo Fisher Scientific) was used to transfer proteins from the gel to a PVDF membrane for long-term storage. Western blots were conducted in accordance with the following protocol as previously described (Agoglia et al. 2017; Faccidomo et al. 2021; Faccidomo et al. 2024b; Wilkie et al. 2007): membranes were incubated in a 3% Normal Goat Serum blocking buffer to minimize non-specific background signal. Next, membranes were incubated overnight at 4°C with one of the following primary antibodies: [rabbit anti-TARP γ-8 (1:8000, S. Tomita, Yale University); mouse anti-CaMKIIα (1:20,000, Millipore #05-532); rabbit antiGluA1(1:1 000, Millipore #04-855); mouse anti-PSD-95 (1:500,000, NeuroMab/Antibodies Inc #75-028); mouse anti-GAPDH (1:10,000; Advanced Immunochemical-1 hr room temperature incubation)]. Membranes were washed and then incubated with a secondary antibody (1:10 000; Jackson ImmunoResearch) and all chemiluminescent signals were digitally imaged (ImageQuant LAS 4000, GE Healthcare Life Sciences) after incubation in ECL Select or Prime (GE Healthcare). The optical density of each band was quantified using ImageQuant TL software and target bands were normalized to the optical density of GAPDH (loading control) and expressed as a ratio (target protein/GAPDH). Within each blot, data are normalized as percent control (sucrose mice).

### Drugs

All oral EtOH (v/v), sucrose (w/v), saccharin (w/v), and quinine (w/v) solutions were prepared by diluting 95% ethanol (Doe and Ingalls), sucrose, saccharin or quinine with tap water. For acute EtOH injection studies, a 20% (w/v) EtOH solution was prepared by diluting 95% ethanol with 0.9% sodium chloride and injection volumes were adjusted to calculate dose.

### Statistical Analysis

Data are presented as MEAN±SEM. All statistical analyses and graphical representations of data were performed using GraphPad Prism software (v. 10.1). Group comparisons were conducted via unpaired t-test, one-way RM ANOVA, or two-way RM ANOVA. Linear regression was used for BAC analyses. Post hoc comparisons were conducted with Dunnett’s or Šídák’s multiple comparison procedures, where appropriate. P values of < 0.05 were considered to be statistically significant. α was set at 0.05 for all comparisons.

## Results

### Appetitive Studies

TARP γ-8 (+/−) mice did not show gross basal differences in body weight, food and water consumption as compared with C57Bl/6J WT mice (**Fig 1B**). For the taste reactivity studies, a two-way mixed ANOVA revealed a significant main effect of concentration on consumption of saccharin (F_(1,16)_=6.45, p=0.022; **Fig 1C**), sucrose (F_(1,16)_=43.84, p<0.0001; **Fig 1C**) and quinine (F_(1,16)_=20.80, p=0.0003; **Fig 1C**). As expected, the higher dose of quinine significantly reduced both intake and preference as compared to water (F_(1,16)_=33.22, p<0.0001) and there was a significant interaction between genotype and concentration for both intake (F_(1,16)_=5.199, p=0.0366; **Fig 1C**) and preference (F_(1,16)_=11.85, p=0.0003; **Fig 1C**). Šídák multiple comparison test revealed that this interaction was due to the differential concentration-dependent quinine intake by TARP γ-8 (+/−) mice (**Fig 1C**).

**Figure 1:**
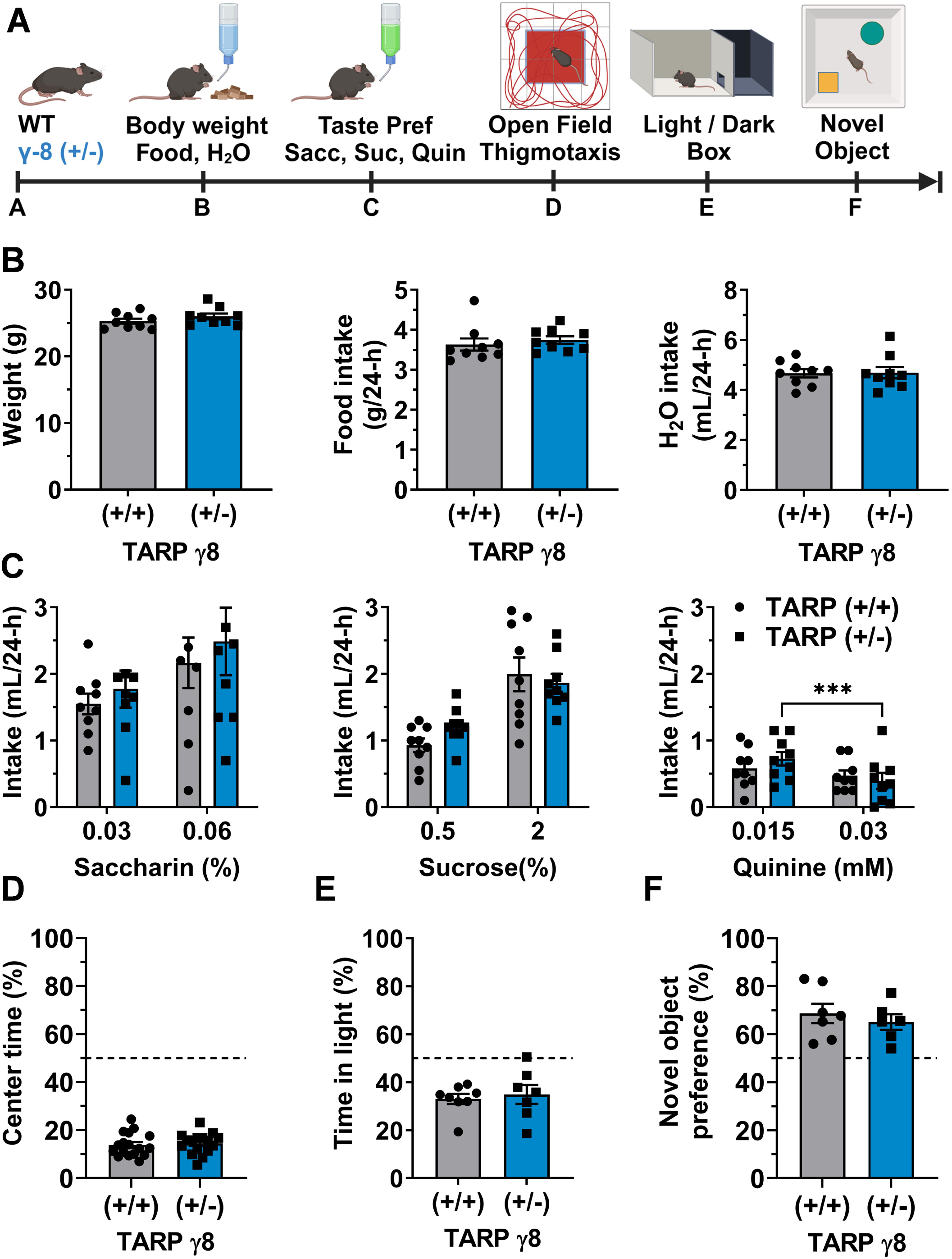
Behavioral characterization of the TARP γ-8 (+/−) mice. Data are expressed as Mean ± SEM, *grey bars* represent TARP γ-8 (+/+) mice and *blue bars* represent TARP γ-8 (+/−) mice. *Asterisks* denote statistical significance (p<0.05) **(A)** Experimental sequence for appetitive, open field, and anxiety-like assessments. **(B)** Comparison of basal body weight (*left panel*), food intake (*center panel*) and water intake (*right panel*), (n=18) showing no genotypic differences. **(C)** Taste reactivity studies (n=18) for two concentrations of Saccharin (*left panel*), Sucrose (*center panel*) and Quinine (*right panel*), *** - significant difference between concentrations, Sidak’s test, P<0.0001. **(D – E)** no genotypic difference in assessments of anxiety-like behavior as measured by time in center of an open field (**D**, n=34) or time in light side of a light/dark chamber (**E**, n=15). (**F**) Novel object recognition test showing preference for novel object irrespective of TARP ɣ-8 genotype (n=13).

### Open Field Locomotion, Anxiety-like Behavior, and Memory

In comparison to C57BL/6J WT mice, TARP γ-8 (+/−) mice did not show statistically significant differences in overall locomotor activity or differences in anxiety-like behavior as measured by time spent in the center zone of an open field (**Fig 1D**), time spent on the light compartment of the light/dark chamber (**Fig 1E**), or time spent with a novel object (**Fig 1F**).

### Acute Ethanol Sensitivity and Blood EtOH Clearance

TARP ɣ-8 (+/−) and (+/+) mice were evaluated for potential differences in response to acute EtOH followed by tests of home-cage EtOH drinking (**Fig 2A**). To assess sensitivity to the acute locomotor activating and sedative hypnotic doses of EtOH, locomotor activity was measured after a 1.0 g/kg dose of EtOH (**Fig 2B**) and loss of righting reflex (**Fig 2C**) was measured after injection of a 4.0 g/kg dose of EtOH. There was not a statistically significant difference between genotypes in either measure. Likewise, both genotypes showed similar blood EtOH clearance curves in response to a 4 g/kg (IP) injection of EtOH (**Fig 2D**).

**Figure 2:**
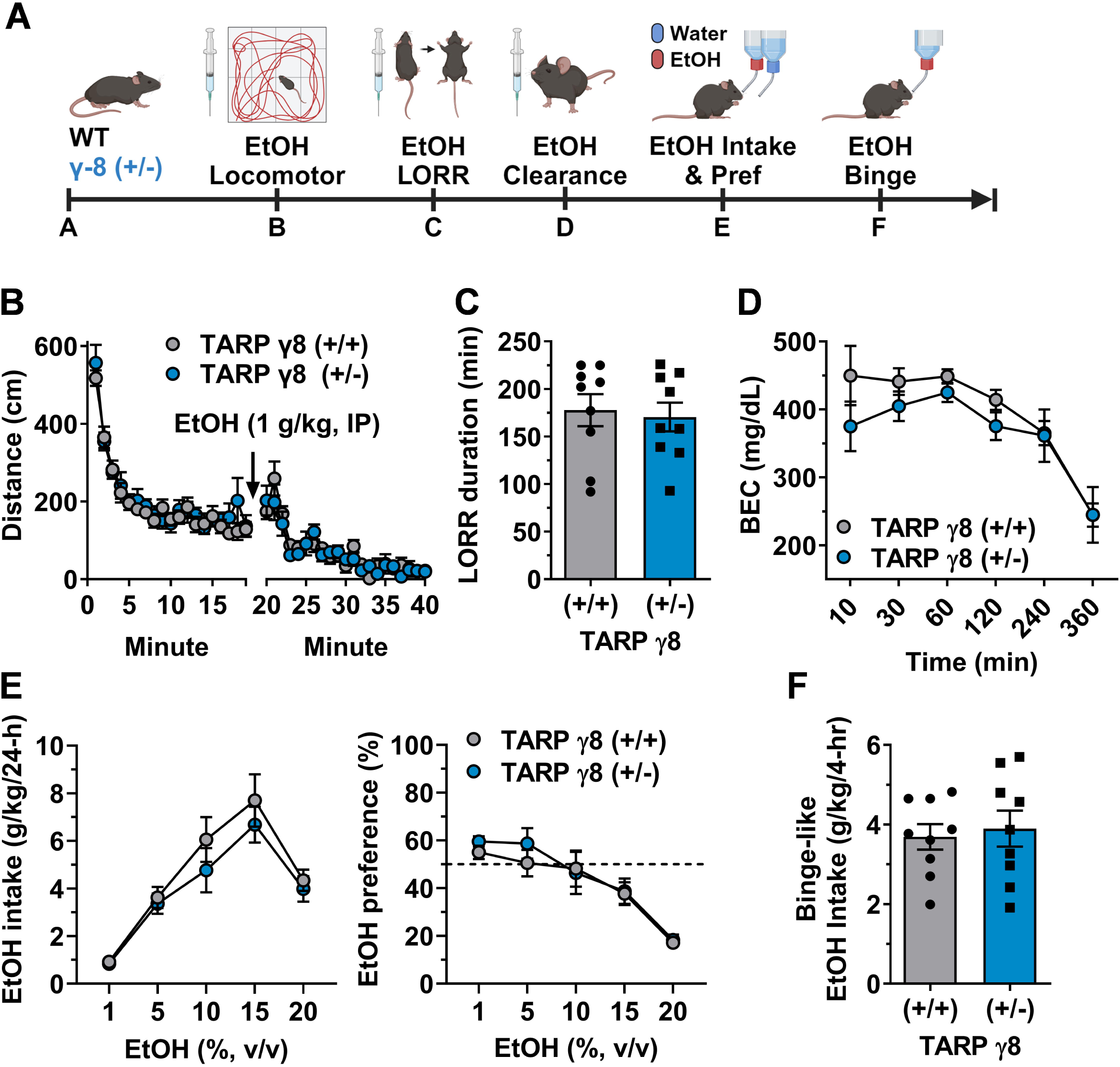
Acute EtOH sensitivity, EtOH clearance and home cage EtOH drinking. Data are expressed as Mean ± SEM, *grey circles and bars* represent TARP γ-8 (+/+) mice and *blue circles and bars* represent TARP γ-8 (+/−) mice. **(A)** Experimental sequence for acute ethanol and home cage EtOH drinking studies. **(B)** No genotypic difference in habituation to the open-field environment or acute locomotor activity following acute EtOH (1 g/kg, IP, n=18) injection. **(C)** No difference in sedation as measured by loss of righting reflex (min) after acute EtOH (4 g/kg, IP, n=18) injection. **(D)** Blood EtOH content (BEC, mg/dL) measured at 6 timepoints after acute EtOH (1 g/kg, IP, n=18) injection. **(E)** No difference in EtOH intake (g/kg/24hrs; *left panel,* and EtOH preference (*right panel*) during continuous-access 2-bottle choice (EtOH vs H_2_O) home-cage drinking test at each of ascending concentration of EtOH. Data are mean of 4 days at each concentration (n=18). **(F)**. No genotypic difference in binge-like EtOH intake (g/kg/4-hr) as tested during limited-access to a single bottle of EtOH (20% v/v) in the home-cage during the dark cycle.

### Home Cage Ethanol Drinking

#### 24-hr access

Mice were given home cage access to water and EtOH, presented in ascending concentrations. Data for each EtOH concentration are expressed as 4-day averages for dose and preference. Although a main effect of genotype was not found, EtOH intake exhibited an inverted U-shaped concentration-response curve in both genotypes with a peak at 15% EtOH (F_(2.196, 35.14)_=38.58, p<0.0001; **Fig 2E**) and post-hoc tests revealed that all EtOH concentrations were significantly increased, compared to intake at 1% EtOH. Further, there was a significant main effect of EtOH concentration on EtOH preference (F_(2.348,37.57)_=29.76, p<0.0001; **Fig 2E**) and post-hoc tests revealed that preference for EtOH vs water was significantly lower at the two highest EtOH concentrations (15-20%).

#### Drinking in the Dark

Both genotypes consumed a comparable amount of 20% (v/v) EtOH during the 4-hr binge session (**Fig 2F**). A microanalysis of the hourly intake during this session did not find any significant differences in the temporal pattern of intake (data not shown).

### Operant Ethanol and Sucrose Self-Administration

#### Acquisition Concentration Curves

To determine if there was an effect of genotype on the acquisition of operant responding for EtOH or sucrose, mice were initially trained to lever press for 2% sucrose only and then tested for potential differences in acquisition and escalation of EtOH self-administration (**Fig 3A**). The TARP γ-8 (+/+) and (+/−) mice showed similar rates of responding for this non-drug reinforcer (**Figure 3B**). Likewise, sucrose control mice that were maintained at 2% sucrose also did not have a main effect of genotype for active lever presses or sucrose intake (data not shown). The data in **Figure 3B** are presented as 3 day means for each EtOH concentration. A 2-way ANOVA (genotype x concentration) revealed a significant main effect of genotype (F _(1, 13)_ = 7.349, p=0.0178), concentration (F_(15, 195)_ = 4.009, p<.0001) and a significant interaction between both factors (F _(15, 195)_ = 2.685, p=0.001). Dunnett’s post-hoc analyses revealed that this interaction was due to an increase in response rate for TARP γ-8 (+/+) mice at 12%, 15% and upon return to 9% EtOH in the reinforcing solution. Specifically, when the reinforcing solution returned to 9% EtOH/2% sucrose, there was a significant escalation in operant rates of responding for EtOH, pre vs post concentration curves that was present in the TARP γ-8 (+/+) mice only for genotype (F _(1, 12)_ = 10.04, p=0.0081), Time (F _(1, 12)_ = 10.19, p=0.0077) and a significant interaction between both factors (F _(1, 12)_ = 7.49, p=0.018; **Figure 3C**). Holm-Šídák’s multiple comparisons test revealed that this interaction was due to increased self-administration by the TARP γ-8 (+/+) mice at the post-time point. A similar result was observed for total number of reinforcements obtained [genotype (F _(1, 12)_ = 10.28, p=0.0076), Time (F _(1, 12)_ = 9.08, p=0.011) and a significant interaction between both factors (F _(1, 12)_ = 7.19, p=0.02); **Figure D**] and dose consumed [g/kg; genotype (F _(1, 12)_ = 7.46, p=0.018), Time (F _(1, 12)_ = 9.71, p=0.009) and a significant interaction between both factors (F _(1, 12)_ = 8.17, p=0.014); **Figure G**]. Significant differences in pre/post timepoints were not seen for % active lever responding or headpoke/reinforcer ratio (**Figure 3E-F**).

**Figure 3:**
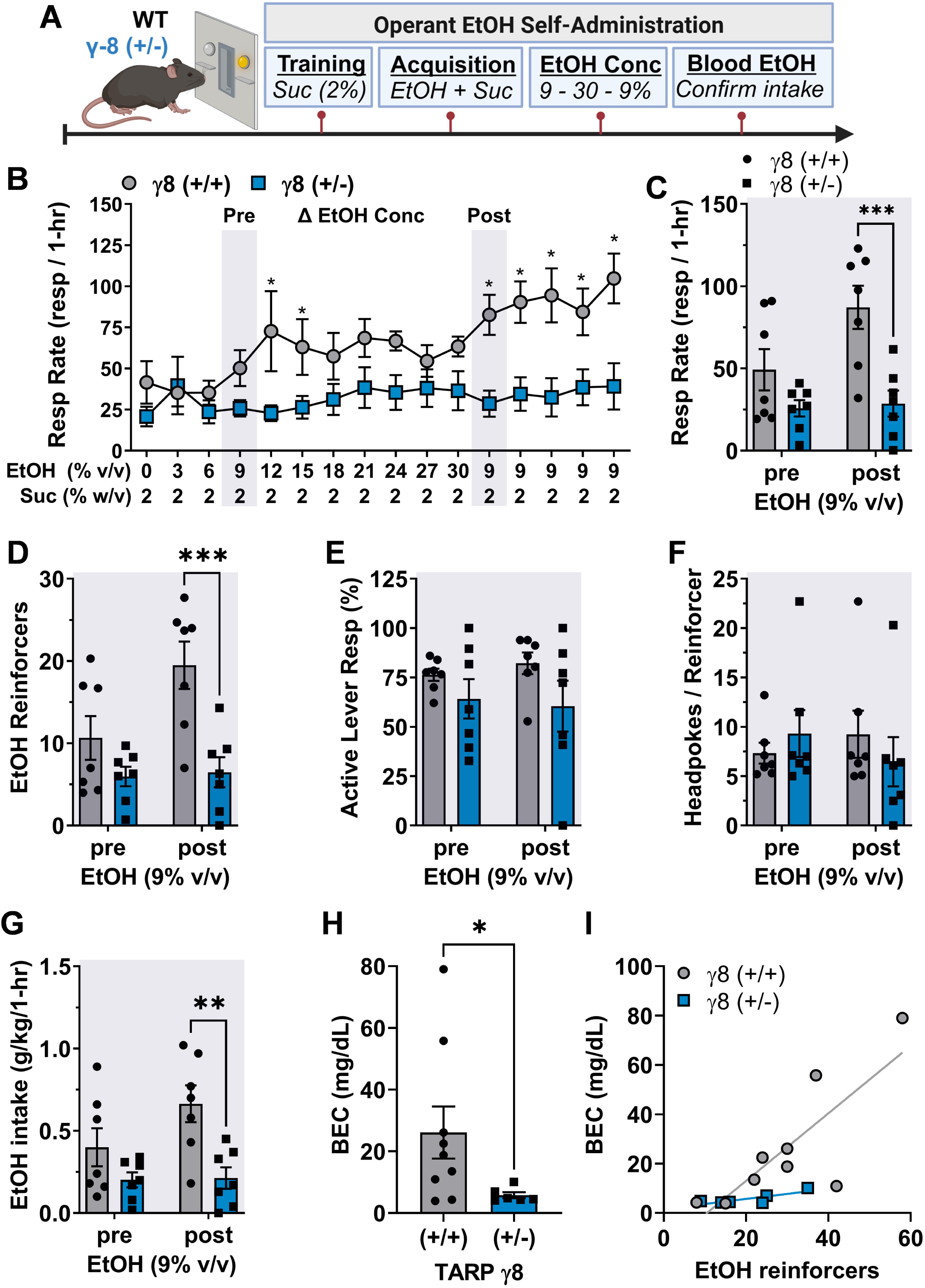
Acquisition and Escalation of Operant Ethanol Self-Administration in TARP γ-8 (+/+) and (+/−) Mice. Data are expressed as Mean ± SEM, with *grey bars/circles* representing TARP γ-8 (+/+) mice (n=8) and *blue bars/circles* representing TARP γ-8 (+/−) mice (n=7). **(A)** Experimental timeline for the acquisition of operant ethanol self-administration following initial training on 2% sucrose. **(B)** Acquisition curves showing the average number of active lever presses over three-day blocks for increasing concentrations of EtOH (2 – 30% v/v) and a return to 9%. **(C)** Escalation of operant alcohol self-administration, showing active lever response rate before and after the escalating concentration protocol (pre vs. post 9% ethanol). **(D)** Total number of reinforcements obtained pre- and post-escalating ethanol concentration. **(E)** Percentage of active lever responding pre- and post-escalating ethanol concentration. **(F)** Headpoke/reinforcer ratio pre- and post-escalating ethanol concentration. **(G)** Dose of ethanol consumed (g/kg) pre- and post-escalating ethanol concentration. **(H)** Blood alcohol content (BAC) levels measured after a 1-hour self-administration session. **(I)** Linear regression analysis showing the relationship between blood alcohol content (BAC) and the number of reinforcements obtained. Regression lines for TARP γ-8 (+/+) and (+/−) mice are shown with *grey and blue lines*, respectively. *Asterisks* denote statistical significance between genotypes, * - P<0.05, Dunnett’s test at each EtOH / Suc concentration **(Panel B)**, ** - p<0.01, *** - p<0.001, Sidak’s test as indicated **(Panels C-D, G)**.

#### Blood EtOH Content

Blood EtOH content (BAC) levels were measured immediately after a 1 hr self-administration session. An unpaired t-test found a significant effect of genotype on BAC levels (t_(13)_ = 1.94, p=0.037; **Figure 3H**). Linear regression was used to interrogate whether there was a relationship between BAC and # of reinforcers obtained. Linear regression analysis of these factors found that the slope of the regression line of TARP γ-8 (+/+) mice was significantly different from 0 (Y = 1.368*X - 14.36; F_(1,7)_=13.26, p=0.008) but the slope of the line for TARP γ-8 (+/−) mice was not (**Figure 3I**).

### Extinction and Cue-Induced Reinstatement

After the acquisition and escalation self-administration concentration curve was conducted, operant responding for 9%EtOH/2% sucrose or 2% sucrose only was maintained for several months until baseline rates of responding were similar for TARP γ-8 (+/+) and TARP γ-8 (+/−) for each reinforcing solution, respectively. After equating baseline performance, mice trained to self-administer EtOH or sucrose underwent extinction followed by three cue-induced reinstatement tests (**Fig 4A and 4D, respectively**)

**Figure 4:**
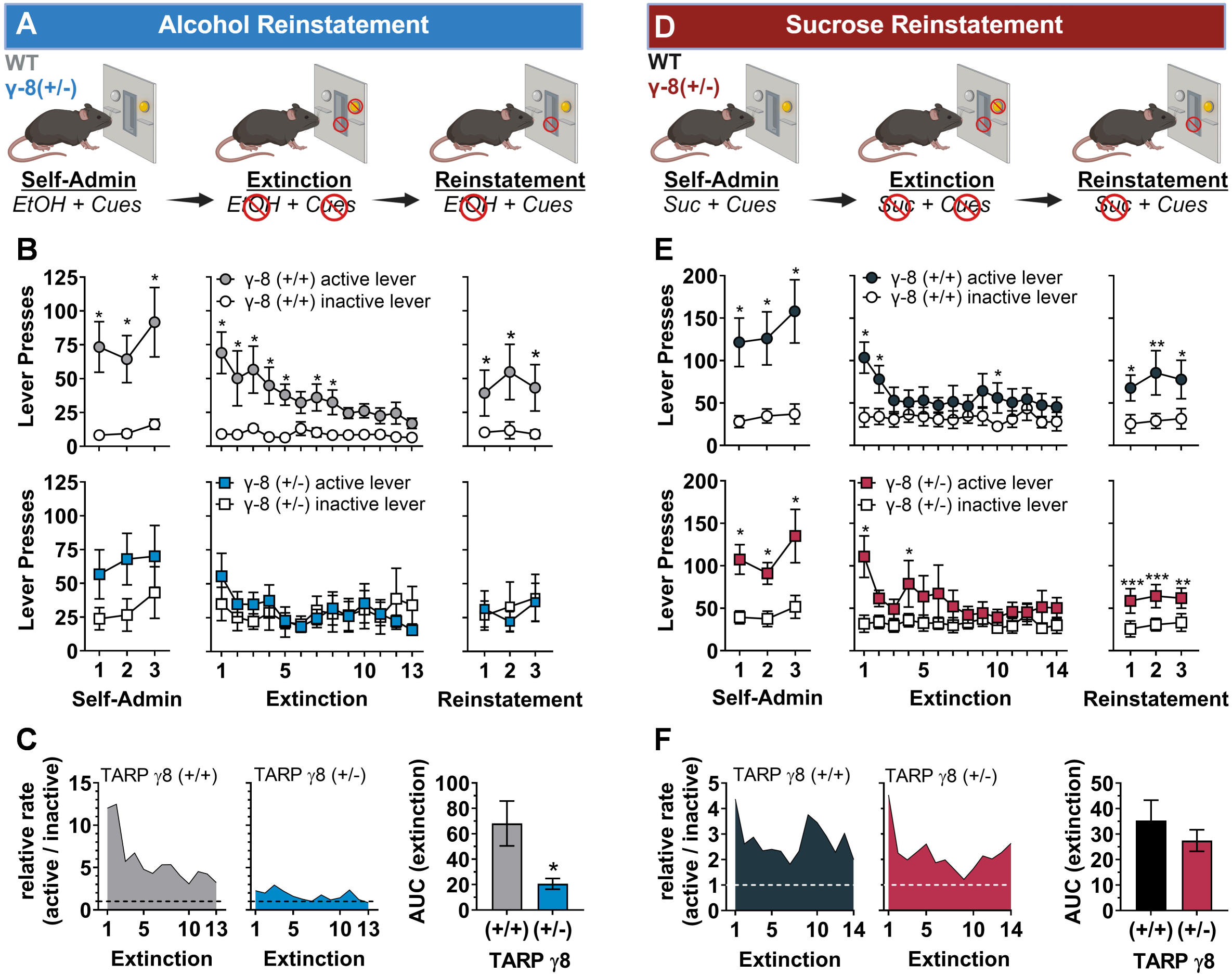
Baseline, Extinction and Cue-Induced Reinstatement of Ethanol and Sucrose Seeking Behavior in TARP γ-8 (+/+) and (+/−) Mice. Data are expressed as Mean ± SEM. *Asterisks* denote statistical significance (p<0.05). **(A and D)** Experimental timeline for baseline, extinction and cue-induced reinstatement following stable self-administration of 9% ethanol/2% sucrose (n=12) or sucrose only (n=17), respectively. **(B, EtOH)** Lever presses on the active and inactive levers during the last three days of FR4 self-administration *(left panels*), during extinction (*center panel*), and during three days of cue-induced reinstatement (*right panels*) for mice trained to self-administer ethanol. **(C, EtOH)** Quantification of the area under the extinction curves for ethanol-trained mice. **(E, Sucrose)** Lever presses on the active and inactive levers during the last three days of FR4 self-administration (*left panels*), during extinction (*center panel*), and during three days of cue-induced reinstatement (*right panels*) for mice trained to self-administer sucrose. **(F, Sucrose)** Quantification of the area under the extinction curves for sucrose-trained mice. *Asterisks* indicate significant differences between genotypes (**Panel C**, unpaired t-test, * - p=0.025) or between active and inactive lever presses (**Panels B and E**, Sidak’s test, * - p<0.05, ** - p<0.01, *** - p<0.001).

#### 9% EtOH/2% sucrose

During the last 3 days of FR4 responding for sweetened EtOH, TARP γ-8 (+/+) mice pressed significantly more times on the active versus inactive lever on all three days (F_(1, 6)_ = 11.76, p=0.014) whereas TARP γ-8 (+/−) (F_(1, 4)_ = 7.1, p=0.056) showed a non-significant trend for increased responding on the active vs inactive lever (**Figure 4B, left panels)**. During extinction, when programmed consequences to responding were eliminated, a 2-way ANOVA revealed that TARP γ-8 (+/+) mice lever pressed significantly more on the active versus inactive lever (F_(1, 6)_ = 8.852, p=0.025) and a significant interaction was found between Extinction Day x Lever (F_(12, 72)_ = 5.547, p<0.0001). Šídák’s multiple comparisons test revealed that active lever responses remained higher than inactive responding for 8 days indicating that operant responding for EtOH was extinguished on the 9^th^ day of extinction for TARP γ-8 (+/+) mice (**Figure 4B, center panel**) TARP γ-8 (+/−) mice did not press significantly more on the active versus inactive lever during extinction indicting rapid extinction to the alcohol -associated cues though there was a main effect of Extinction Day (F_(12,48)_ = 2.316, p=0.012; **Figure 4B, center panel**). Post-hoc tests revealed that responding on both levels was significantly reduced by day 5 vs day 1 of extinction. Quantification of the area under the extinction curves (t_(10)_=2.176, p=0.026; **Figure 4C**) revealed that extinction rates between TARP γ-8 (+/+) and TARP γ-8 (+/−) mice significantly differed. To assess cue-induced EtOH reinstatement, mice were placed into the operant conditioning chamber and visual, olfactory and auditory secondary cues associated with the reinforcer delivery were presented, but reinforcers were not delivered. Reinstatement tests were repeated for 3 consecutive days. TARP γ-8 (+/+) (F_(1, 6)_ = 6.107, p=0.085) but not TARP γ-8 (+/−) mice responded significantly more on the active versus inactive lever across all 3 reinstatement sessions (**Figure 4B, right panels)**.

#### 2% sucrose

During the last 3 days of FR4 responding for EtOH, both TARP γ-8 (+/+) (F_(1, 7)_ = 10.51, p=0.014) and TARP γ-8 (+/−) F_(1, 8)_ = 16.39, p=0.004) mice pressed significantly more times on the active versus inactive lever on all three days (**Figure 4E, left panels**). In the sucrose-trained TARP γ-8 (+/+) mice, a 2-way ANOVA revealed a significant interaction was found between Extinction Day x Lever (F_(13, 91)_ = 2.424, p=0.007; **Figure 4E, center panel**). Šídák’s multiple comparisons test revealed that active lever responses remained significantly higher than inactive responding on days 1, 2 and 10 of extinction. Likewise, in sucrose-trained TARP γ-8 (+/−) mice, a 2-way ANOVA revealed that these mice lever pressed significantly more on the active versus inactive lever (F_(1, 8)_ = 6.90, p=0.030) and a significant interaction was found between Extinction Day x Lever (F_(13, 104)_ = 2.066, p=0.022; **Figure 4E, center panel**). Šídák’s multiple comparisons test revealed that active lever responses remained significantly higher than inactive responding on days 1 and 4 of extinction indicating that the TARP γ-8 (+/−) mice extinguished active lever responding faster than the TARP γ-8 (+/+) mice. Quantification of the area under the extinction curves (t_(11)_=2.321, p=0.045; **Figure 4F**) revealed that extinction rates between TARP γ-8 (+/+) and TARP γ-8 (+/−) mice did not differ by genotype. To assess cue-induced sucrose reinstatement, mice were placed into the operant conditioning chamber and visual, olfactory and auditory secondary cues associated with the reinforcer delivery were presented, but reinforcers were not delivered. Reinstatement tests were repeated for 3 consecutive days. TARP γ-8 (+/+) (F_(1, 7)_ =4.74, p=0.0659) showed a variable but strong trend for repeated reinstatement, whereas but TARP γ-8 (+/−) (F_(1, 8)_ =6.81, p=0.031) mice showed full reinstatement of sucrose-seeking behavior across all 3 reinstatement sessions, but planned comparisons within each day showed that both genotypes responded more on the previously active sucrose lever as compared to the inactive lever (**Figure 4E, right panels)**.

### Protein Expression

To address whether a history of sucrose or EtOH operant self-administration altered protein expression throughout the brain, levels of protein immunoreactivity for TARP γ-8, GluA1, CamKIIα and PSD-95 were compared within genotype between sucrose- and EtOH -trained mice for 4 key brain regions. There were no significant differences between genotype for CaMKIIα and PSD-95 in any brain region (data not shown); hence, protein changes for TARP γ-8 and GluA1 are reported below.

#### Amygdala

TARP γ-8 (+/+) mice had a significantly increased level of both TARP γ-8 immunoreactivity (t_(12)_=4.636, p=0.0006) and GluA1 (t_(12)_=4.46, p=0.0008) immunoreactivity in EtOH vs sucrose-trained mice (**Figure 5A, black and grey bars, left panels**). TARP γ-8 (+/−) mice did not show increased TARP γ-8 immunoreactivity but did have a significantly increased level of GluA1 (t_(11)_=4.986, p=0.0004) immunoreactivity in EtOH vs sucrose-trained mice (**Figure 5A, red and blue bars, right panels**).

**Figure 5:**
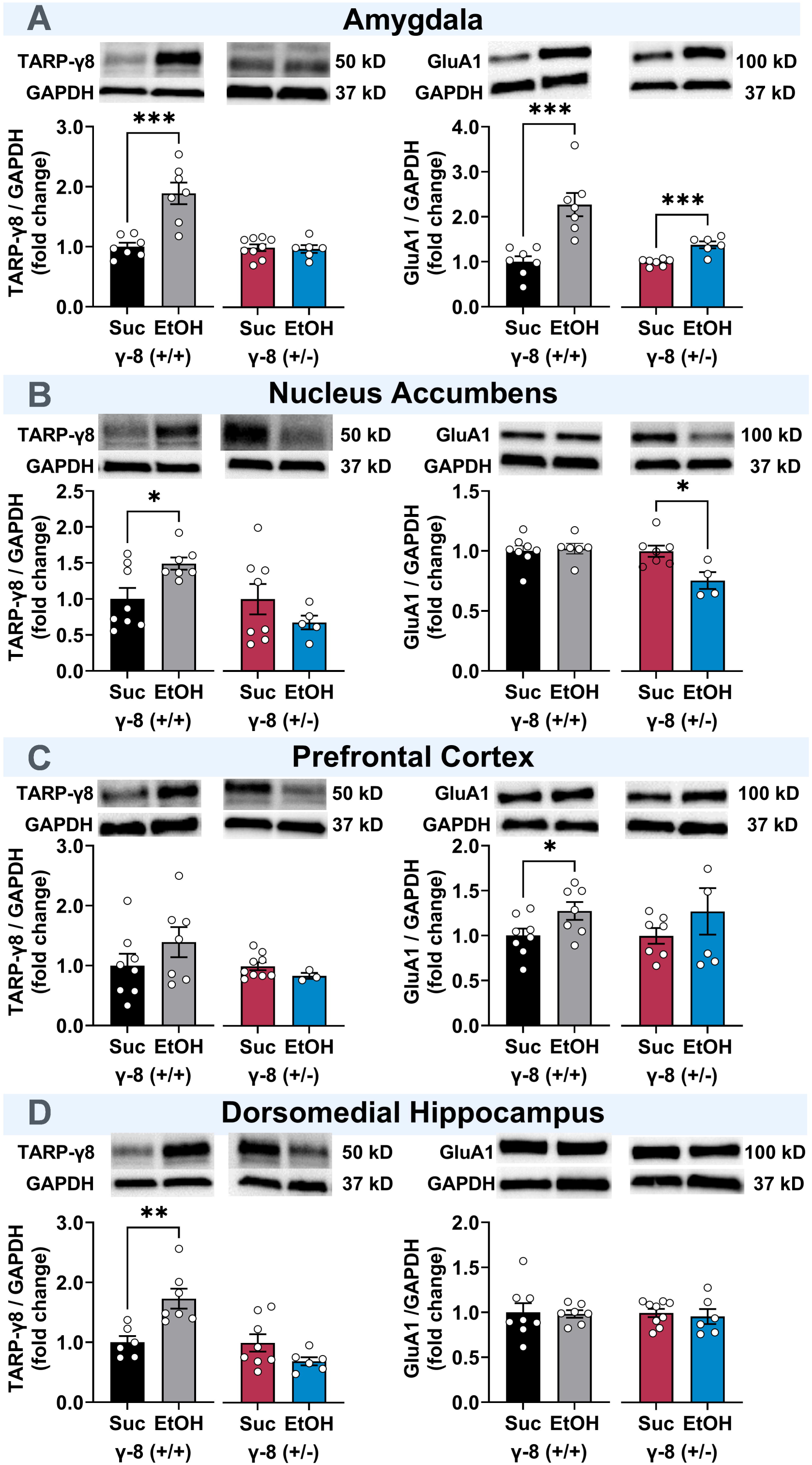
Brain Region-Specific Changes in TARP γ-8 and GluA1 Protein Expression Following Operant Ethanol Self-Administration. Protein immunoreactivity levels for TARP γ-8 and total GluA1 (GluA1) were compared between ethanol self-administering (n=13) and sucrose self-administering (n=16) mice for TARP γ-8 (+/+) and TARP γ-8 (+/−) mice. Data are presented as Mean ± SEM. *Asterisks* denote statistical significance (unpaired t-test, * - p<0.05, ** - p<0.01, *** - p<0.001) **(A) Amygdala:** *Left panels* show TARP γ-8 (+/+) mice (Ethanol: *black bars*, Sucrose: *grey bars*) exhibited significantly increased levels of both TARP γ-8 and GluA1 immunoreactivity in ethanol-trained compared to sucrose-trained mice. *Right panels* show TARP γ-8 (+/−) mice (Ethanol: *red bars*, Sucrose: *blue bars*) did not show increased TARP γ-8 immunoreactivity but had significantly increased GluA1 immunoreactivity in ethanol-trained mice. **(B) Nucleus Accumbens:** *Left panels* show TARP γ-8 (+/+) mice (Ethanol: *black bars*, Sucrose: *grey bars*) had significantly increased TARP γ-8 immunoreactivity but no change in GluA1 immunoreactivity in ethanol-trained mice. *Right panels* show TARP γ-8 (+/−) mice (Ethanol: *red bars*, Sucrose: *blue bars*) did not show increased TARP γ-8 immunoreactivity but had significantly decreased GluA1 immunoreactivity in ethanol-trained mice. **(C) Medial Prefrontal Cortex:** *Left panels* show TARP γ-8 (+/+) mice (Ethanol: *black bars*, Sucrose: *grey bars*) showed a trend towards increased TARP γ-8 but a significant increase in GluA1 immunoreactivity in ethanol-trained mice. *Right panels* show TARP γ-8 (+/−) mice (Ethanol: *red bars*, Sucrose: *blue bars*) showed no change in immunoreactivity for either protein. **(D) Dorsomedial Hippocampus:** *Left panels* show TARP γ-8 (+/+) mice (Ethanol: *black bars*, Sucrose: *grey bars*) had significantly increased TARP γ-8 immunoreactivity but no change in GluA1 immunoreactivity in ethanol-trained mice. *Right panels* show TARP γ-8 (+/−) mice (Ethanol: *red bars*, Sucrose: *blue bars*) showed no change in immunoreactivity for either protein.

#### Nucleus Accumbens

TARP γ-8 (+/+) mice had a significantly increased level of TARP γ-8 immunoreactivity (t_(13)_=2.689, p=0.019) but no change in GluA1 immunoreactivity in EtOH vs sucrose-trained mice (**Figure 5B, black and grey bars, left panels**). In contrast, TARP γ-8 (+/) mice did not show increased TARP γ-8 immunoreactivity but did have a significantly decreased level of GluA1 immunoreactivity (t_(9)_=2.982, p=0.015) in EtOH vs sucrose-trained mice (**Figure 5B, red and blue bars, right panels**).

#### Medial Prefrontal Cortex

TARP γ-8 (+/+) mice trended towards but did not show a significantly increased level of TARP γ-8 immunoreactivity in the mPFC but did show a significant increase in GluA1 (t_(12)_=2.184, p=0.0479) immunoreactivity in EtOH vs sucrose-trained mice (**Figure 5C, black and grey bars, left panels**). TARP γ-8 (+/−) mice did not show a change in immunoreactivity for either protein in the mPFC (**Figure 5C, red and blue bars, right panels**).

#### Dorsomedial Hippocampus

TARP γ-8 (+/+) mice had a significantly increased level of TARP γ-8 immunoreactivity (t_(11)_=3.571, p=0.004) but no change in GluA1 immunoreactivity in EtOH vs sucrose-trained mice (**Figure 5D, black and grey bars, left panels**). In contrast, TARP γ-8 (+/) mice did not show increased TARP γ-8 or GluA1 immunoreactivity in EtOH vs sucrose-trained mice (**Figure 5D, red and blue bars, right panels**).

## Discussion

This study establishes TARP γ-8 as a critical molecular determinant of EtOH reinforcement and relapse vulnerability. By demonstrating that partial genetic deletion of TARP γ-8 selectively attenuates both the reinforcing effects of EtOH and cue-induced relapse-like behavior, we provide novel insights into the underlying molecular mechanisms of alcohol misuse. These findings strongly suggest that TARP γ-8 represents a promising target for the development of therapeutic interventions for AUD.

### Basal Functions

To determine if TARP γ-8 (+/−) mice were free of health or behavioral anomalies that could confound the evaluation of EtOH-related behaviors, we assessed several general basal parameters. The absence of differences in body weight, food consumption, and water intake between TARP γ-8 (+/−) mice and wildtype controls suggests that partial reduction of TARP γ-8 expression does not impact general metabolic or homeostatic functions. Additionally, the lack of differences in taste preference for saccharin, sucrose, or quinine indicate that the reinforcing effects of EtOH observed in this study are not driven by alterations in gustatory perception in TARP γ-8 (+/−) mice (e.g., (Hodge et al. 1999)). Similar performance by TARP ɣ-8 (+/−) mice and wildtype controls in the open-field (thigmotaxis) and the light-dark box further suggests that differences in EtOH self-administration are not attributable to differences in baseline anxiety, which can be predictive of altered EtOH intake in mouse models (Hodge et al. 2004; Kelley et al. 2003). Finally, the absence of any deficits in memory performance in the novel object recognition task indicates that TARP γ-8 (+/−) mice do not exhibit global impairments in learning or memory; however, this test does not directly assess associative learning or memory, which is regulated by AMPAR activity (Rumpel et al. 2005). Taken together, these null findings suggest that the behavioral alterations observed in EtOH reinforcement and relapse-like behavior in TARP γ-8 (+/−) mice are specific to the neurobiological mechanisms underlying EtOH reinforcement and are not due to broader changes in metabolism, anxiety, or cognitive function.

### Acute Ethanol Response

TARP γ-8 (+/−) mice were evaluated for potential differences in acute ethanol responses that might influence ethanol self-administration (Hodge et al. 2004; Hodge et al. 1999). Locomotor activity assessments revealed no differences between TARP γ-8 (+/−) mice and wildtype controls in habituation to a novel environment, indicating that exploratory behavior and motor function are intact. Furthermore, both genotypes exhibited comparable motor activation and subsequent habituation following acute EtOH (1 g/kg, IP) administration, suggesting that TARP γ-8 does not play a role in modulating adaptation to a novel environment or the initial locomotor stimulant effects of EtOH. When examining the sedative-hypnotic effects of a higher dose of EtOH (4 g/kg, IP), no differences were observed in the duration of loss of righting reflex (LORR) between TARP γ-8 (+/−) mice and wildtype controls, indicating similar sensitivity to the depressant effects of EtOH. Finally, blood ethanol analysis showed no differences in absorption or clearance rates, suggesting that EtOH pharmacokinetics are not altered by partial TARP γ-8 reduction. These findings collectively suggest that the observed differences in EtOH self-administration and relapse-like behavior in TARP γ-8 (+/−) mice are not attributable to changes in EtOH metabolism or acute motor responses, further supporting the specificity of TARP γ-8 in modulating ethanol’s reinforcing properties.

### Ethanol Reinforcement: Requirement of TARP ɣ-8

The reduction in the acquisition of operant EtOH self-administration observed in TARP γ-8 heterozygous null mice (+/−) compared to wild-type controls underscores the potential role of TARP γ-8 in the development of AUD. Notably, this reduction was specific to EtOH reinforcement. TARP γ-8 (+/−) mice showed no difference in sucrose-reinforced responding during acquisition suggesting the absence of a learning deficit and underscoring potential specificity of regulation of EtOH reinforcement. Interestingly, TARP ɣ-8 (+/−) mice exhibited no differences from wild-type controls in voluntary EtOH consumption when assessed via home-cage 24-hour two-bottle choice drinking (across a range of EtOH concentrations from 1 to 20% v/v) or in 4-hour binge-like access sessions to a single bottle containing 20% EtOH. These findings suggest that the role of TARP γ-8 in EtOH self-administration is not due to altered EtOH preference or intake under free-access conditions but is more likely tied to the reinforcement mechanisms driving operant EtOH self-administration.

Furthermore, wild-type mice exhibited a two-fold increase in operant EtOH (9% v/v) self-administration following exposure to an escalating EtOH concentration protocol (12 - 30% v/v), a model of increasing EtOH intake often associated with the development of AUD. In contrast, TARP γ-8 (+/−) mice did not show this concentration-induced escalation in EtOH self-administration, indicating that TARP γ-8 is essential for the dynamic reinforcement processes that promote increased EtOH seeking and consumption under higher EtOH exposure conditions. The absence of this escalation in TARP γ-8 (+/−) mice highlights a potential protective effect of reduced TARP γ-8 expression against the transition from controlled EtOH use to problematic or escalated drinking, which characterizes AUD.

This selectivity indicates that TARP γ-8 plays a targeted role in modulating the reinforcing properties of EtOH, rather than affecting general reward processing or behaviors unrelated to EtOH reinforcement. Importantly, TARP γ-8 (+/−) mice did not display any differences in motor function or EtOH metabolism, further supporting the specificity of TARP γ-8’s role in the neural circuits that govern EtOH reinforcement and escalation. This is consistent with our previous pharmacological studies demonstrating that the functions of TARP γ-8-bound AMPARs are selectively involved in EtOH reinforcement, with no impact on non-drug reinforcement (e.g., sucrose), motor activity, or anxiety-like behavior (Hoffman et al. 2021; Hoffman et al. 2023).

These findings support the conclusion that TARP γ-8 is essential in modulating the development of EtOH’s reinforcing properties and underscores the potential importance of TARP γ-8 in the neuroadaptations that contribute to escalated EtOH use that characterizes the progression of AUD. This suggests that TARP γ-8 may serve as a valuable target for therapeutic interventions aimed at preventing or reducing escalated drinking and the progression of AUD.

### Role of TARP γ-8 in Relapse Vulnerability

In addition to playing a role in EtOH reinforcement, results of this study show that TARP γ-8 also appears to be a critical regulator of extinction of conditioned reinforcement and cue-induced relapse-like behavior, with specificity to EtOH. Our findings demonstrate that TARP γ-8 (+/−) mice exhibit rapid extinction and significantly diminished cue-induced reinstatement of ethanol-seeking behavior across three repeated cue-exposure tests; however, extinction of sucrose-reinforced responding and reinstatement were unaltered by TARP ɣ-8 deletion. This specific of behavioral function suggests that TARP γ-8 serves a key mechanistic role in regulating neural processes that underlie relapse to alcohol-seeking behavior.

Our previous research has shown that cue-induced reinstatement of ethanol-seeking behavior is associated with elevated pCaMKII-T286 immunoreactivity in several reward-related brain regions, including the amygdala, nucleus accumbens, and piriform cortex (Salling et al. 2017). This suggests that CaMKII signaling is engaged in response to EtOH-associated cues, potentially driving synaptic plasticity changes critical for relapse. When considered in conjunction with the present findings, it is plausible that the absence or reduction of TARP γ-8 may disrupt this CaMKII-AMPAR signaling cascade, leading to a reduction in synaptic insertion of TARP γ-8-bound AMPARs in response to EtOH-related cues. Consequently, this could attenuate cue-induced reinstatement of EtOH-seeking behavior. Future investigations should directly evaluate this hypothesis by quantifying synaptic AMPAR levels and phosphorylated CaMKII in TARP γ-8 (+/−) mice during cue-induced reinstatement, which could provide critical evidence for the proposed mechanism.

Another significant finding from this study is that TARP γ-8 (+/−) mice exhibited immediate extinction of EtOH-seeking behavior, indicating a profound reduction in behavioral momentum, or persistence, of the discriminated operant response (Nevin 2012). According to behavioral momentum theory, the persistence of operant behavior during extinction is directly related to the history of reinforcement, particularly its frequency and density. The lower levels of EtOH self-administration observed in TARP γ-8 (+/−) mice during acquisition and extended baseline phases resulted in a diminished reinforcer density, making their ethanol-seeking behavior more susceptible to extinction when ethanol was no longer available. By contrast, the persistence of ethanol-seeking behavior in wild-type mice suggests that TARP γ-8 regulation of AMPAR-mediated synaptic transmission may enhance ethanol reinforcement and strengthen resistance to extinction (Cannady et al. 2013; Cannady et al. 2017), suggesting that modulating TARP γ-8 activity could be a strategy to weaken the persistence of drug-seeking behavior. Future studies are warranted to determine how TARP γ-8 influences the persistence of chronic EtOH-seeking behavior; however, these studies will require equating baseline exposure of TARP ɣ-8 null mice or using pharmacological tools, such as JNJ-55511118 to block TARP ɣ-8 bound AMPARS only during extinction.

A particularly compelling finding of this study is the ethanol-specificity of the rapid extinction observed in TARP γ-8 (+/−) mice, since they exhibited the same rate of extinction for sucrose-reinforced responding as wild-type controls. This specificity is consistent with prior research demonstrating that inhibition of TARP γ-8-bound AMPARs selectively reduces the reinforcing effects of ethanol while having no impact on sucrose reinforcement (Hoffman et al. 2021). These findings suggest that TARP γ-8 regulation of AMPARs may be particularly important in modulating extinction and persistence of drug-seeking behaviors, rather than only affecting EtOH reinforcement. Future studies should explore whether TARP γ-8 similarly regulates the extinction and reinstatement of other drugs of abuse, which could provide broader insights into its role in addiction-related neural plasticity.

The rapid extinction and absence of cue-induced reinstatement of EtOH-seeking behavior in TARP γ-8 (+/−) mice raise important questions about potential deficits in associative learning or memory (Park et al. 2016). The reduced reinforcing function of EtOH in these mice likely led to weaker associations between the conditioned stimulus (CS, environmental cues linked to EtOH) and EtOH’s unconditioned stimulus (US) function. Consequently, this diminished association could reduce the likelihood of these cues eliciting EtOH-seeking behavior during extinction or reinstatement. This weak association may stem from reduced availability of extrasynaptic TARP γ-8-bound AMPARs (AMPARs located outside of the synapse, which play a role in synaptic plasticity) and impaired modulation of AMPAR activity, which are crucial for the excitatory synaptic transmission needed to establish and recall learned associations (Kato et al. 2016; Kato et al. 2010a; Kato et al. 2010b; Rumpel et al. 2005; Sumioka et al. 2011). In TARP γ-8 null mice, impairments in associating environmental cues with EtOH may reduce their ability to respond effectively to these cues, suggesting deficits in AMPAR-dependent encoding and retrieval of cue-EtOH associations, which are critical for behaviors involved in addiction (Hyman 2005; Hyman et al. 2006; Kelley 2004), highlighting the importance of TARP γ-8 in these fundamental learning processes.

Overall, our findings suggest that modulation of AMPARs by TARP γ-8 influences both the persistence of EtOH-seeking behavior (extinction) and cue-induced relapse, suggesting that targeting TARP γ-8 could offer a novel therapeutic approach to addressing the neurobiological mechanisms of addiction and reducing relapse vulnerability.

### Protein Expression: Impact of ethanol on TARP γ-8 and AMPAR GluA1

A novel finding of this study is that operant EtOH self-administration significantly increased TARP γ-8 protein expression in the amygdala, nucleus accumbens, and dorsal hippocampus, but not prefrontal cortex, in wildtype control mice as compared to sucrose self-administration. However, this effect was absent in TARP γ-8 (+/−) mice. This finding underscores the anatomical specificity of EtOH effects on TARP γ-8 expression as compared to a non-drug reinforcer, suggesting that TARP γ-8 plays a crucial role in modulating specific brain regions and neural circuits underlying EtOH reinforcement.

Notably, we also observed that EtOH self-administration increased AMPA receptor GluA1 expression in the amygdala in wildtype mice, as compared to sucrose. Interestingly, while GluA1 expression also increased in the amygdala of TARP γ-8 (+/−) mice following EtOH self-administration compared to sucrose, this increase was noticeably blunted compared to the robust increase observed in wildtype mice. This suggests that TARP γ-8 contributes to the full extent of EtOH-induced GluA1 upregulation in this key reward-related brain region. These results highlight the unique pathways activated by EtOH compared to the non-drug reinforcer sucrose and provide insights into the molecular underpinnings of AUD. Understanding how TARP γ-8 modulates effects of EtOH, including its influence on GluA1 expression, may lead to targeted interventions for EtOH use disorders, as auxiliary proteins like TARP γ-8 are critical for AMPA receptor function and synaptic plasticity, which are pivotal in addiction neurobiology.

The absence of changes in PSD-95 protein expression in the context of altered TARP γ-8 and GluA1 expression suggests a specific modulation of glutamate receptor signaling pathways that does not involve alterations in the postsynaptic density scaffold proteins. PSD-95 is a key scaffolding protein in the postsynaptic density that organizes AMPA receptors and other signaling molecules, and it plays a critical role in synaptic plasticity and function (Hodge et al. 2004) and EtOH -induced changes in synaptic structure (Mulholland and Chandler 2007) and behavior (Camp et al. 2011). In our results, the increase in TARP γ-8 and GluA1 protein expression suggests enhanced trafficking or stabilization of AMPA receptors at the synapse, potentially leading to heightened synaptic efficacy. However, the lack of change in PSD-95 suggests that the underlying synaptic architecture remains intact, despite the alterations in receptor composition induced by EtOH self-administration. It is possible that TARP γ-8 and GluA1 are participating in dynamic regulation of AMPA receptor activity independent of PSD-95 associated structural influence.

### Potential Compensatory Changes

A crucial consideration in using null mouse models is the potential for compensatory changes in the absence of the targeted gene(s). Initial characterization of TARP γ-8 heterozygous (γ-8+/−) mice showed that there were no differences in synaptic transmission between γ-8+/− and wild-type mice, indicating that heterozygous mice maintain normal AMPAR and NMDAR function (Rouach et al. 2005). This suggests that a partial deletion of TARP γ-8 does not impair synaptic AMPAR-mediated transmission or alter synaptic strength, as evidenced by unchanged AMPA/NMDA ratios, EPSP slope, paired-pulse facilitation, and mEPSC frequency. While γ-8+/− mice show small reductions in AMPAR GluA1 and GluA2/3 subunit expression, these changes do not affect the receptor composition, as the rectification index (RI) remained unchanged between genotypes (Rouach et al. 2005). The preserved synaptic function and receptor composition in γ-8+/− mice suggest that any observed effects in the present study are due to TARP γ-8 function rather than compensatory changes in AMPAR systems. However, a complete understanding of the role of TARP ɣ-8 in behavioral response to EtOH will require further study of potential compensatory changes in other proteins and use of additional approaches to manipulate TARP ɣ-8 activity.

### Conclusion and Implications for Therapeutic Development

This study establishes TARP γ-8 as a critical molecular mechanism underlying core behavioral pathologies associated with AUD: positive reinforcement, escalated consumption, extinction, and cue-induced relapse, with no effects in parallel sucrose controls. Importantly, TARP γ-8 exhibits restricted anatomical expression in key limbic regions implicated in EtOH-related behaviors, such as the prefrontal cortex, hippocampus, and basolateral amygdala, all of which are integral components of reward circuitry involving excitatory projections to the nucleus accumbens. This specific localization strongly suggests that targeting TARP ɣ-8 has potential to transform AUD treatment by offering a precise intervention in AMPAR signaling within restricted reward-related neural circuits, with potentially fewer side effects than broad-acting therapies. Indeed, coupled with our prior demonstration that the selective TARP γ-8-associated AMPAR inhibitor JNJ-55511118 effectively reduces EtOH self-administration in preclinical models (Hoffman et al. 2021; Hoffman et al. 2023), these compelling findings firmly position pharmacological modulation of TARP γ-8 as a novel and promising therapeutic strategy in the medical management of AUD. It is important to note that these findings were obtained using an animal model. Thus, although these preclinical results offer strong support for the role of TARP γ-8 in AUD, the translation of these findings to human patients with AUD remains an empirical question that warrants further investigation.

## Acknowledgement

Funding for this research was provided by grants R01 AA028782 (CWH) and P60 AA011605 (CWH and MAH) and by support from the Bowles Center for Alcohol Studies at UNC Chapel Hill.

Timeline images in Figures 1 – 3 (Panel A) and Figure 4 (Panels A and D) were created with BioRender.com by CWH.

## Author contributions

SF, JLH, ST, AEG, MAH, and CWH developed the conceptual and methodological bases for this work. JL, CMW, MK, SMT, AK, CR, HLS, JMB, AC, and ENS collected and organized data. SF, JLH, and CWH analyzed and graphed data. CWH and SF drafted the manuscript. All authors contributed to study designs, interpretation of results, and approved the final version of the manuscript.

## Data availability

Data sets used in this study are available upon reasonable request.

## Ethics declarations

## Ethnical approval

All procedures were performed in accordance with institutional guidelines and according to the NIH Guide for the Care and Use of Laboratory Animals (NIH Publication 80-23) and with the Institutional Animal Care and Use Committee at the University of North Carolina at Chapel Hill.

